# First evidence for TIGR-Tas activity in a photosynthetic eukaryote

**DOI:** 10.64898/2026.09.12.750307

**Authors:** Jaeyeon Lee, Kyeongjun Kim, Hyeonmin Ryu, Hyeonsik Yoon, Kerstin Dürr, Mid-Eum Park, Joong-Tak Yoon, Bon-Kyoung Koo, Ji-Hye Yoon, Ho-Seok Lee, Eun Yu Kim

**Author notes:** Correspondence: Jaeyeon Lee, Ho-Seok Lee, Eun Yu Kim.

## Abstract

TIGR–Tas systems are a recently described class of RNA-guided DNA-targeting systems whose activity has been demonstrated in bacterial and human cells, but not in photosynthetic eukaryotes. Here we tested TIGR–Tas in the green alga *Chlamydomonas reinhardtii*. Recombinant TaTas and ParTas assembled with in vitro-transcribed tigRNAs cleaved target DNA in a concentration-dependent manner, and electroporated TaTas protein was detected in *C. reinhardtii* cells. Targeted deep sequencing of the endogenous *MAA7* locus recovered indel-containing reads in ParTas-treated cells that were absent from wild-type controls, including a 1-bp deletion located at the position previously reported for ParTas-associated cleavage. Co-delivery of a double-stranded non-homologous oligonucleotide (dsNHO) with a 24-bp duplex and 8-nt 3′ overhangs, designed to match the staggered ends generated by TasR-family nucleases, increased the recovery of indel-containing reads for both proteins. The frequency of these events was very low, and the present data do not establish TIGR–Tas as an efficient genome-editing system in this host. They nonetheless provide initial evidence that TIGR–Tas can act on the nuclear genome of a photosynthetic eukaryote, and indicate that dsNHO-assisted recovery of sequence alterations is not restricted to conventional CRISPR– Cas nucleases.

## Introduction

*Chlamydomonas reinhardtii* (*C. reinhardtii*) is a well-established model photosynthetic eukaryote that combines a genetically tractable haploid genome, rapid growth, and simple cultivation with extensive molecular and genomic resources.[1] These features have made *C. reinhardtii* an important system for studying photosynthesis, organelle biology, and cellular metabolism, while also supporting its development as a biotechnology platform.[2] Genome engineering has further expanded its experimental utility, with CRISPR–Cas systems enabling targeted gene disruption and sequence modification.[3] However, only a limited range of programmable nuclease systems has been experimentally validated across microalgae.[1] Expanding this repertoire with mechanistically distinct programmable nucleases is valuable not because such systems necessarily replace CRISPR–Cas, but because alternative targeting architectures may provide complementary properties and broaden the range of molecular strategies available for genome manipulation. *C. reinhardtii* therefore provides a useful photosynthetic context in which to evaluate the functionality and host compatibility of newly discovered genome-engineering systems.[1]

TIGR–Tas systems, recently identified as a distinct class of RNA-guided DNA-targeting systems, use tandem-spacer guide RNAs to direct Tas proteins to DNA through a targeting architecture fundamentally different from conventional CRISPR–Cas systems.[4] Nuclease-active Tas proteins can mediate programmable DNA cleavage and have demonstrated genome-editing activity in heterologous cells, but their functionality in photosynthetic eukaryotes has not yet been established.[4] Testing TIGR–Tas in *C. reinhardtii* therefore provides an opportunity to determine whether this recently discovered RNA-guided system can operate in a phylogenetically and cellularly distinct host. Because the activity of newly introduced nucleases can also depend strongly on host DNA-repair responses, strategies that enhance the recovery of editing events may be particularly important when initial activity is low. Here, we evaluated TIGR–Tas-mediated genome modification in *C. reinhardtii* and examined whether co-delivery of a non-homologous oligonucleotide could increase the recovery of TIGR–Tas-associated sequence alterations.[5]

## Results

To establish an RNP-based TIGR–Tas genome-editing system in *C. reinhardtii*, we first purified TaTas and ParTas proteins for subsequent biochemical and cellular experiments. Both Tas proteins contained a nuclear localization signal (NLS) and His tag and had a calculated molecular weight of approximately 45.6 kDa. SpCas9 containing an NLS and His tag, with a calculated molecular weight of 161.5 kDa, was also purified for the in vitro cleavage assay. SDS–PAGE analysis showed prominent protein bands corresponding to the expected molecular sizes of TaTas, ParTas, and SpCas9, confirming successful preparation of the proteins (Figure 1A).

**Figure 1.**
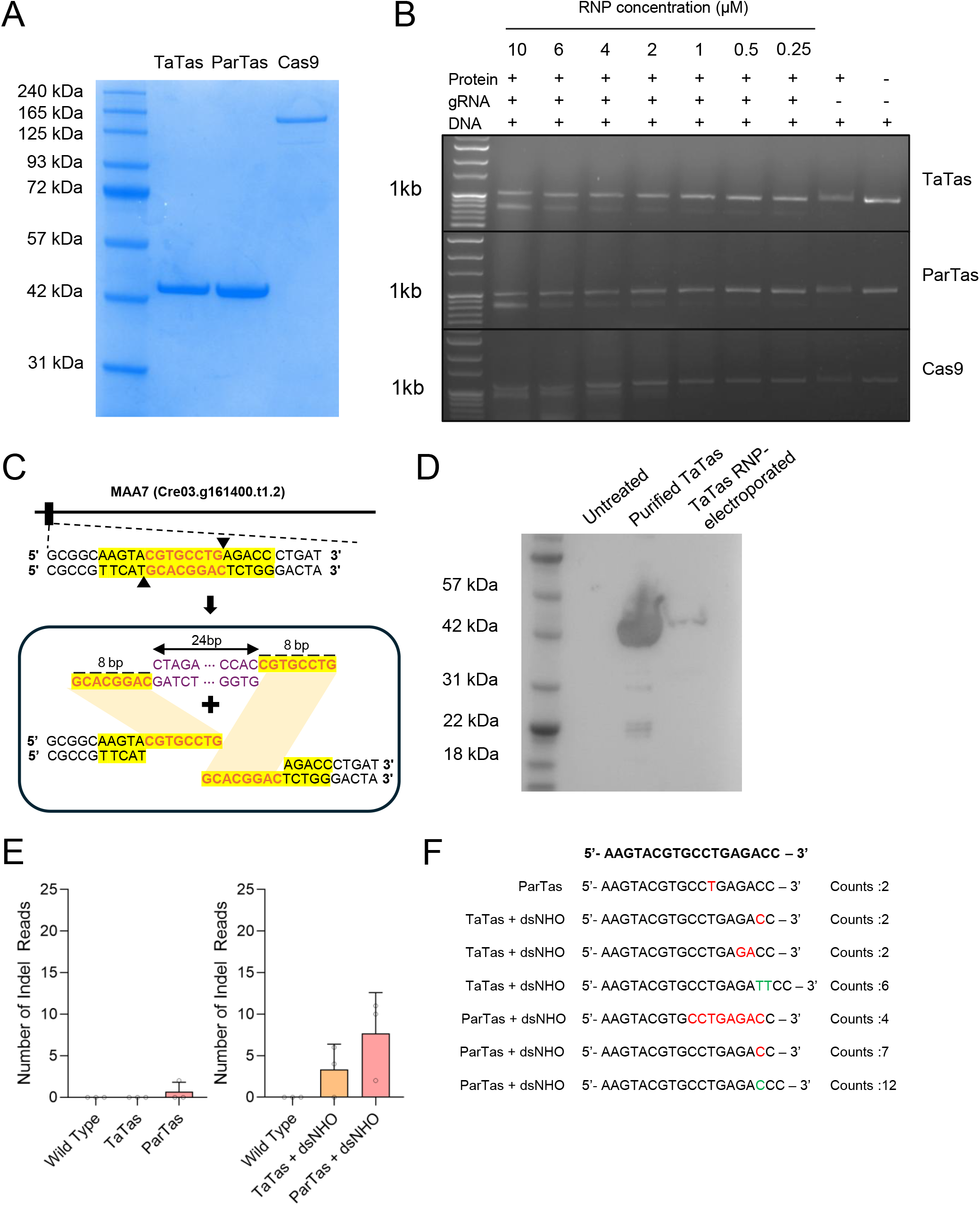
RNP-based TIGR–Tas targeting of the MAA7 locus in Chlamydomonas reinhardtii. (**A**) SDS–PAGE analysis of purified TaTas, ParTas, and SpCas9 proteins. (**B**) In vitro cleavage assay using TaTas, ParTas, and SpCas9 RNPs assembled with in vitro-transcribed tigRNA or sgRNA. (**C**) Schematic of the *MAA7* target region and the double-stranded non-homologous oligonucleotide (dsNHO) used for co-delivery with TIGR–Tas RNPs. (**D**) Western blot detection of His-tagged TaTas following electroporation into *Chlamydomonas reinhardtii* CC-4533 cells. Purified recombinant TaTas was loaded as a positive control; untreated CC-4533 cells served as a negative control. (**E**) Targeted deep-sequencing analysis of the *MAA7* locus following TaTas or ParTas RNP delivery with or without dsNHO. The y-axis indicates the number of indel-containing reads detected by deep sequencing. (**F**) Representative mutant sequences identified by deep sequencing, with the wild-type reference sequence shown at the top. Deletions are indicated in red and insertions in green; counts indicate the number of corresponding reads.

We next examined whether the purified Tas proteins retained programmable DNA-cleavage activity. tigRNAs for TaTas and ParTas, together with sgRNA for SpCas9, were prepared by in vitro transcription and assembled with their corresponding purified proteins to form ribonucleoprotein (RNP) complexes. The resulting RNPs were incubated with the target DNA at 37°C for 20 min over a concentration range of 0.25–10 μM (Figure 1B). Both TaTas and ParTas cleaved the 1-kb target DNA, generating approximately 800-bp and 200-bp DNA fragments, with cleavage becoming more evident as the RNP concentration increased. SpCas9 RNP also cleaved its corresponding target DNA under the tested conditions, producing the expected cleavage fragments. In contrast, the protein-only and DNA-only controls did not show comparable cleavage products. These results confirmed that the purified TaTas and ParTas proteins retained concentration-dependent DNA-cleavage activity following assembly with their respective tigRNAs. We subsequently investigated whether TIGR–Tas could induce genome modification in *C. reinhardtii*.

The endogenous *MAA7* gene (Cre03.g161400.t1.2) was selected as the genomic target (Figure 1C). In addition to delivery of TIGR–Tas RNP alone, we examined whether co-delivery of a double-stranded non-homologous oligonucleotide (dsNHO) could enhance the recovery of TIGR–Tas-mediated editing events. Previous work in *C. reinhardtii* demonstrated that dsNHO co-delivery can enhance CRISPR-mediated gene knockout and that this phenomenon is influenced by dsNHO length, structure, and chemical modification and is largely dependent on the KU70/80 DNA double-strand break sensor.[5] It further proposed that dsNHOs interfere with cellular DSB sensing and alter the balance between canonical non-homologous end joining and microhomology-mediated end joining.[5] Because the reported enhancement was observed across different Cas nucleases, genomic loci, and strains, we asked whether a similar strategy could be applied to TIGR–Tas, a distinct RNA-guided nuclease system that generates staggered DNA ends.[5] In contrast to the previous study, in which dsNHOs containing 4-nt 5′ overhangs were designed based on the staggered cleavage pattern of Cas12a, we adapted the dsNHO design to the cleavage characteristics of TIGR– Tas.[5] TasR-family nucleases, including TaTas and ParTas, generate staggered DNA cleavage with an 8-nt offset between the two strands, producing 8-nt 3′ overhangs.[4] Accordingly, we designed a dsNHO consisting of 32-nt oligonucleotides that form a 24-bp duplex with 8-nt 3′ overhangs at both ends and co-delivered it with preassembled TIGR–Tas RNPs during electroporation (Figure 1C).

Before evaluating genome-editing outcomes, we established conditions for TIGR–Tas RNP delivery into the *C. reinhardtii* CC-4533 strain using TaTas. Preassembled TaTas RNPs were introduced into CC-4533 cells by electroporation, and intracellular TaTas protein was subsequently examined by western blotting using an anti-His antibody (Figure 1D). No corresponding signal was detected in untreated wild-type cells, whereas a protein band corresponding to TaTas was detected specifically in the electroporated samples. This result confirmed that TaTas protein could be successfully delivered into CC-4533 cells under the electroporation conditions used for the following genome-editing experiments.

Deep-sequencing analysis of the targeted *MAA7* region was performed in three independent experiments for each treatment condition (Figure 1E). Because the frequency of altered reads was extremely low, the results are presented as the number of reads containing indels rather than as a conventional editing efficiency. The wild-type control libraries contained 20,640, 59,649, and 48,760 total reads, respectively, and no indel-containing reads were detected. Similarly, no indel-containing reads were detected in any of the three TaTas-only libraries, which contained 20,934, 14,003, and 13,625 total reads, respectively. In contrast, two deletion-containing reads were identified across the three ParTas-only libraries, which contained 20,342, 20,482, and 23,073 total reads, respectively.

Co-delivery of the dsNHO increased the recovery of altered reads for both nucleases. Four deletion-containing reads were detected across the three TaTas plus dsNHO libraries, which contained 31,698, 21,258, and 25,537 total reads. The strongest signal was observed for ParTas plus dsNHO, for which 23 indel-containing reads were recovered from libraries containing 30,665, 28,000, and 33,370 total reads. Thus, although the absolute number of altered reads remained very low relative to the total sequencing depth, dsNHO co-delivery was associated with increased recovery of TIGR–Tas-related sequence alterations, particularly in the ParTas condition (Figure 1E).

Representative sequences of indel-containing reads are shown in Figure 1F. ParTas alone generated low-frequency deletion-containing reads, whereas dsNHO co-delivery increased the recovery of indel-containing reads for both TaTas and ParTas. Notably, a larger deletion was observed in the ParTas plus dsNHO condition, whereas the other recovered variants predominantly consisted of short indels. The 1-bp deletion detected in the ParTas condition occurred at the same relative position with respect to the predicted cleavage site as a deletion previously reported during the initial characterization of the TIGR–Tas system. Based on the reported top-strand cleavage position of ParTas, the deleted thymine was located 2 bp from the cleavage site, consistent with the previously observed ParTas-associated deletion pattern (Figure 1F). Although the frequency of this event was very low, the positional correspondence between the recovered deletion and the previously characterized ParTas cleavage-associated mutation provides additional support that the observed sequence alteration was associated with TIGR–Tas activity rather than a randomly distributed sequence variant.

In summary, we tested the TIGR–Tas system in the photosynthetic eukaryote *C. reinhardtii*. Both TaTas and ParTas showed concentration-dependent DNA-cleavage activity in vitro, and TIGR–Tas-associated indels were recovered following RNP delivery into CC-4533, although at very low frequency. The 1-bp deletion recovered in the ParTas condition occurred at the position expected from the previously characterized ParTas cleavage site, supporting an association with TIGR–Tas activity. Co-delivery of a dsNHO designed to match the staggered ends generated by TasR-family nucleases increased the recovery of indel-containing reads for both proteins, including in the TaTas condition, in which RNP delivery alone yielded no altered reads. This suggests that dsNHO-assisted enhancement may not be restricted to conventional CRISPR–Cas nucleases.

Several limitations should be considered when interpreting these findings. First, the frequency of indel-containing reads was extremely low, and the present data therefore do not establish TIGR–Tas as an efficient genome-editing system in *C. reinhardtii*. Second, only a single endogenous locus and one *C. reinhardtii* strain were examined, and additional targets and genetic backgrounds will be required to determine the generality of TIGR–Tas activity in this host. Third, although dsNHO co-delivery was associated with increased recovery of altered reads, the limited number of editing events precludes a definitive assessment of the magnitude or statistical robustness of this effect. Within these limitations, the present data provide, to our knowledge, initial evidence for TIGR–Tas activity in a photosynthetic organism, and support further investigation with additional target loci, independent biological replicates, and optimized delivery and repair conditions.

## Material and Method

### Ribonucleoprotein (RNP) Formation

Expression vectors encoding Tas and Cas proteins were transformed into *Escherichia coli* BL21 competent cells. Starter cultures were grown in Terrific Broth (TB) supplemented with 50 μg/mL kanamycin or 100 μg/mL ampicillin for 12 h. The starter cultures were subsequently used to inoculate 1 L of TB, and the cells were cultured at 37°C until the OD_600_ reached approximately 0.6. Protein expression was induced by the addition of IPTG to a final concentration of 0.5 mM, followed by incubation at 16°C for 16 h.

Cells were harvested by centrifugation at 4,000 × *g* for 20 min at 4°C. The cell pellets were resuspended in lysis buffer containing 50 mM Tris-HCl (pH 7.5), 500 mM NaCl, and protease inhibitor cocktail. Cells were disrupted by sonication, and the lysates were clarified by centrifugation at 15,000 × *g* for 30 min at 4°C.

The clarified lysates were applied to Ni-NTA resin. The resin was washed with wash buffer containing 25 mM Tris-HCl (pH 7.5), 1 M NaCl, and 30 mM imidazole. Bound proteins were subsequently eluted with elution buffer containing 50 mM Tris-HCl (pH 7.5), 500 mM NaCl, 300 mM imidazole, and protease inhibitor cocktail. The purified proteins were concentrated using Amicon Ultra centrifugal filters, and protein concentrations were determined using a NanoDrop spectrophotometer.

sgRNAs and tigRNAs were synthesized by in vitro transcription using the Guide-it™ sgRNA In Vitro Transcription Kit (Takara Bio) according to the manufacturer’s instructions. For RNP formation, the purified proteins were mixed with their corresponding sgRNAs or tigRNAs and incubated at 37°C for 20 min.

### Invitro cleavage assay

In vitro cleavage assays were performed as previously described.[4]

### Electroporation

*C. reinhardtii* CC-4533 cells were cultured and prepared at a density of 1 × 10^8^ cells/mL. The cells were washed twice with MAX Efficiency™ Transformation Reagent. The prepared cells were mixed with the RNP complexes and transferred to 2-mm-gap electroporation cuvettes. Electroporation was performed using a Gene Pulser Xcell™ Electroporation System (Bio-Rad, USA).

### Targeted deep sequencing

Genomic regions encompassing the target sites were amplified by PCR, and the resulting amplicons were subjected to targeted amplicon deep sequencing on an Illumina MiSeq platform using 100-bp paired-end sequencing (2 × 100 bp). Sequence reads were analyzed to determine the frequencies of insertions and deletions (indels) at the target sites.

## Author Contributions

J.L. contributed to the study design, performed the experiments, and wrote the manuscript. K.K. contributed to manuscript writing. H.Y., M.P., and J.Y. contributed to the study design. H.R. contributed to the protein purification experiments. K.D. contributed to the in vitro cleavage assays. B.K.K. contributed to the supervision of the study. J.H.Y. supervised the protein purification experiments. H.S.L. and E.Y.K. contributed to funding acquisition, study design, and manuscript writing.

## Data availability

Raw sequencing data will be deposited in the NCBI SRA and made available upon publication.

## Acknowledgements

Special Thanks to Dr. Trang Thi Le and Dr. Yong Jae Lee. This research was supported by grants from the Institute for Basic Science (IBS-R021-D1-2026-a00) to H.-S.L. and E.Y.K., the National Research Foundation of Korea (RS-2024-00338015) to H.-S.L., the Research Fund for International Scientists (RFIS), National Natural Science Foundation of China (NSFC), China (32350610246), and the Kunshan Science and Technology Bureau, China (KSSC202302072) to E.Y.K.. No conflict of interest is declared.

